# A systematic approach for dissecting the molecular mechanisms of transcriptional regulation in bacteria

**DOI:** 10.1101/239335

**Authors:** Nathan M. Belliveau, Stephanie L. Barnes, William T. Ireland, Daniel L. Jones, Mike J. Sweredoski, Annie Moradian, Sonja Hess, Justin B. Kinney, Rob Phillips

## Abstract

Gene regulation is one of the most ubiquitous processes in biology. But while the catalog of bacterial genomes continues to expand rapidly, we remain ignorant about how almost all of the genes in these genomes are regulated. At present, characterizing the molecular mechanisms by which individual regulatory sequences operate requires focused efforts using low-throughput methods. Here we show how a combination of massively parallel reporter assays, mass spectrometry, and information-theoretic modeling can be used to dissect bacterial promoters in a systematic and scalable way. We demonstrate this method on both well-studied and previously uncharacterized promoters in the enteric bacterium *Escherichia coli.* In all cases we recover nucleotide-resolution models of promoter mechanism. For some promoters, including previously unannotated ones, the approach allowed us to further extract quantitative biophysical models describing input-output relationships. This method opens up the possibility of exhaustively dissecting the mechanisms of promoter function in *E. coli* and a wide range of other bacteria.

---

The sequencing revolution has left in its wake an enormous challenge: the rapidly expanding catalog of sequenced genomes is far outpacing a sequence-level understanding of how the genes in these genomes are regulated. This ignorance extends from viruses to bacteria to archaea to eukaryotes. Even in *E. coli,* the model organism in which transcriptional regulation is best understood, we still have no indication if or how more than half of the genes are regulated (Fig. S1; see also RegulonDB (1) or EcoCyc (2)). In other model bacteria such as *Bacillus subtilis, Caulobacter crescentus, Vibrio harveyii,* or *Pseudomonas aeruginosa,* far fewer genes have established regulatory mechanisms (3–5).

New approaches are needed for studying regulatory architecture in these and other bacteria. Although an arsenal of genetic and biochemical methods have been developed for dissecting promoter function at individual bacterial promoters (reviewed in Minchin *et al.* (6)), these methods are not readily parallelized. As a result, they will likely not lead to a comprehensive understanding of full regulatory genomes anytime soon. RNA sequencing, chromatin immunoprecipitation, and other high-throughput techniques are increasingly being used to study gene regulation in *E. coli* (7–11), but these methods are incapable of revealing either the nucleotide-resolution location of all functional transcription factor binding sites, or the way in which interactions between DNA-bound transcription factors and RNA polymerase modulate transcription.

In recent years a variety of massively parallel reporter assays have been developed for dissecting the functional architecture of transcriptional regulatory sequences in bacteria, yeast, and metazoans. These technologies have been used to infer biophysical models of well-studied loci, to characterize synthetic promoters constructed from known binding sites, and to search for new transcriptional regulatory sequences (12–18). CRISPR assays have also shown promise for identifying longer range enhancer-promoter interactions in mammalian cells (19). However, no approach for using massively parallel reporter technologies to decipher the functional mechanisms of previously uncharacterized regulatory sequences has yet been established.

Here we describe a systematic and scalable approach for dissecting the functional architecture of previously uncharacterized bacterial promoters at nucleotide resolution using a combination of genetic, functional, and biochemical measurements. First, a massively parallel reporter assay (Sort-Seq (12)) is performed on a promoter in multiple growth conditions in order to identify functional transcription factor binding sites. DNA affinity chromatography and mass spectrometry (20, 21) are then used to identify the regulatory proteins that recognize these sites. In this way one is able to identify both the functional transcription factor binding sites and cognate transcription factors in previously unstudied promoters. Subsequent massively parallel assays are then performed in gene-deletion strains to provide additional validation of the identified regulators. The reporter data thus generated is also used to infer sequence-dependent quantitative models of transcriptional regulation. In what follows, we first illustrate the overarching logic of our approach through application to four previously annotated promoters: *lacZYA, relBE, marRAB,* and *yebG.* We then apply this strategy to the previously uncharacterized promoters of *purT, xylE,* and *dgoRKADT,* demonstrating the ability to go from complete regulatory ignorance to explicit quantitative models of a promoter’s input-output behavior.

## Results

To dissect how a promoter is regulated, we begin by performing Sort-Seq (12). As shown in Fig. 1A, Sort-Seq works by first generating a library of cells, each of which contains a mutated promoter that drives expression of GFP from a low copy plasmid (5-10 copies per cell (22)) and provides a read-out of transcriptional state. We use fluorescence-activated cell sorting (FACS) to sort cells into multiple bins gated by their fluorescence level and then sequence the mutated plasmids from each bin. We found it sufficient to sort the libraries into four bins and generated data sets of about 0.5-2 million sequences across the sorted bins (Fig. S3A-D). To identify putative binding sites, we calculate ‘expression shift’ plots that show the average change in fluorescence when each position of the regulatory DNA is mutated (Fig. 1B, top plot). Mutations to the DNA will in general disrupt binding of transcription factors (23), so regions with a positive shift are suggestive of binding by a repressor, while a negative shift suggests binding by an activator or RNA polymerase (RNAP).

**Fig. 1.**
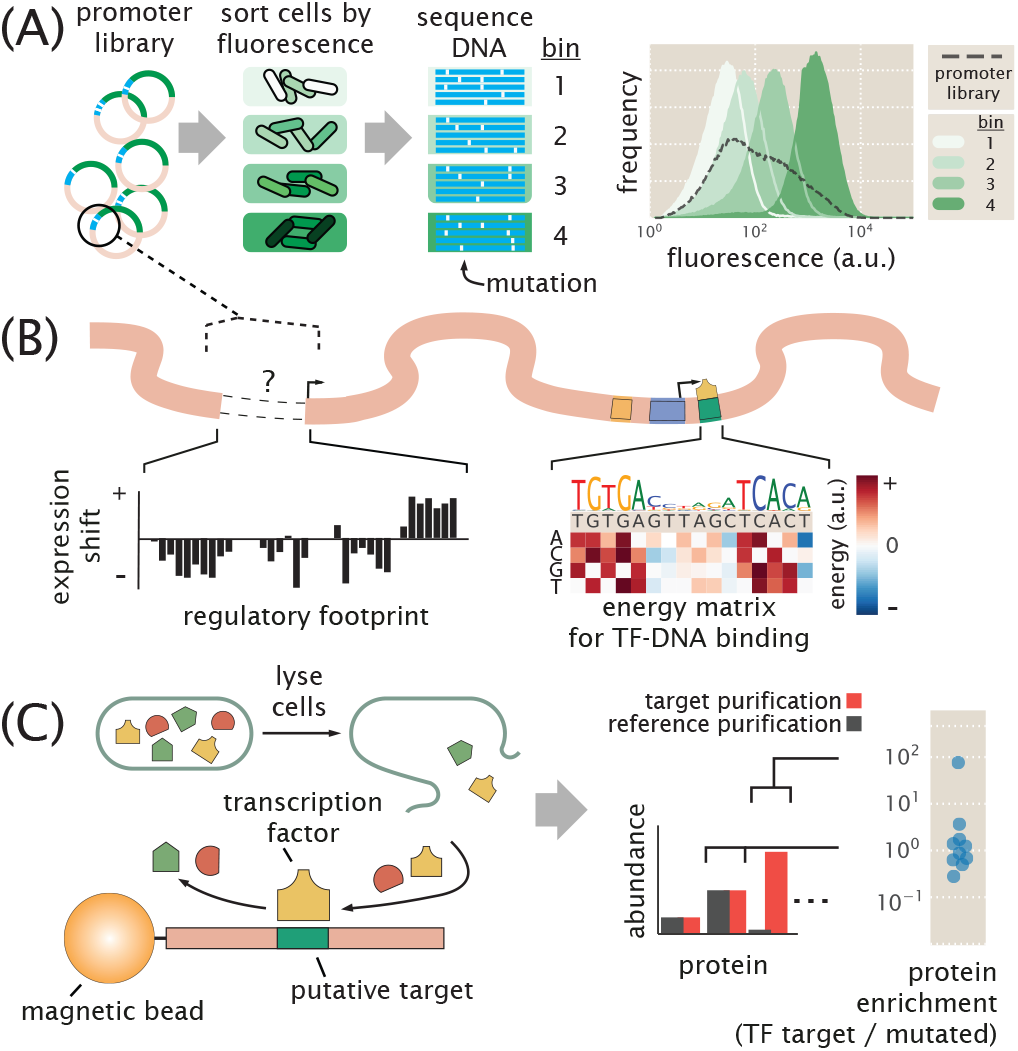
Overview of approach to characterize transcriptional regulatory DNA, using Sort-Seq and mass spectrometry. (A) Schematic of Sort-Seq. A promoter plasmid library is placed upstream of GFP and is transformed into cells. The cells are sorted into four bins by FACS and after regrowth, plasmids are purified and sequenced. The entire intergenic region associated with a promoter is included on the plasmid and a separate downstream ribosomal binding site sequence is used for translation of the *GFP* gene. The fluorescence histograms show the fluorescence from a library of the *rel* promoter and the resulting sorted bins. (B) Regulatory binding sites are identified by calculating the average expression shift due to mutation at each position. In the schematic, positive expression shifts are suggestive of binding by repressors, while negative shifts would suggest binding by an activator or RNAP. Quantitative models can be inferred to describe the associated DNA-protein interactions. An example energy matrix that describes the binding energy between an as yet unknown transcription factor to the DNA is shown. By convention, the wild-type nucleotides have zero energy, with blue squares identifying mutations that enhance binding (negative energy), and where red squares reduce binding (positive energy). The wild-type sequence is written above the matrix. (C) DNA affinity chromatography and mass spectrometry is used to identify the putative transcription factor (TF) for an identified repressor site. DNA oligonucleotides containing the target binding site are tethered to magnetic beads and used to purify the target transcription factor from cell lysate. Protein abundance is determined by mass spectrometry and a protein enrichment is calculated as the ratio in abundance relative to a second reference experiment where the target sequence is mutated away.

The identified binding sites are further interrogated by performing information-based modeling with the Sort-Seq data. Here we generate energy matrix models (12, 24) that describe the sequence-dependent energy of interaction of a transcription factor at each putative binding site. For each matrix, we use a convention that the wild-type sequence is set to have an energy of zero (see example energy matrix in Fig. 1B). Mutations that enhance binding are identified in blue, while mutations that weaken binding are identified in red. We also use these energy matrices to generate sequence logos (25) which provides a useful visualization of the sequence-specificity (see above matrix in Fig. 1B).

In order to identify the putative transcription factors, we next perform DNA affinity chromatography experiments using DNA oligonucleotides containing the binding sites identified by Sort-Seq. Here we apply a stable isotopic labeling of cell culture (SILAC (26)) approach, which enables us to perform a second reference affinity chromatography that is simultaneously analyzed by mass spectrometry. We perform chromatography using magnetic beads with tethered oligonucleotides containing the putative binding site (Fig. 1C). Our reference purification is performed identically, except that the binding site has been mutated away. The abundance of each protein is determined by mass spectrometry and used to calculate protein enrichment ratios, with the target transcription factor expected to exhibit a ratio greater than one. The reference purification ensures that non-specifically bound proteins will have a protein enrichment near one. This mass spectrometry data and the energy matrix models provide insight into the identity of each regulatory factor and potential regulatory mechanisms. In certain instances these insights then allow us to probe the Sort-Seq data further through additional information-based modeling using thermodynamic models of gene regulation. As further validation of binding by an identified regulator, we also perform Sort-Seq experiments in gene deletion strains, which should no longer show the associated positive or negative shift in expression at their binding site.

### Sort-Seq recovers the regulatory features of well-characterized promoters

To first demonstrate Sort-Seq as a tool to discover regulatory binding sites *de novo* we began by looking at the promoters of *lacZYA* (*lac*), *relBE* (*rel*), and *marRAB* (*mar*). These promoters have been studied extensively (27–29) and provide a useful testbed of distinct regulatory motifs. To proceed we constructed libraries for each promoter by mutating their known regulatory binding sites. (See Supplemental Information Section B and Fig. S3E,F for additional characterization). We begin by considering the *lac* promoter, which contains three *lac* repressor (LacI) binding sites, two of which we consider here, and a cyclic AMP receptor (CRP) binding site. It exhibits the classic catabolic switch-like behavior that results in diauxie when *E. coli* is grown in the presence of glucose and lactose sugars (27). Here we performed Sort-Seq with cells grown in M9 minimal media with 0.5% glucose. The expression shifts at each nucleotide position are shown in Fig. 2A, with annotated binding sites noted above the plot. The expression shifts reflect the expected regulatory role of each binding site, showing positive shifts for LacI and negative shifts for CRP and RNAP. The difference in magnitude at the two LacI binding sites likely reflect the different binding energies between these two binding site sequences, with LacI O3 having an *in vivo* dissociation constant that is almost three orders of magnitude weaker than the LacI O1 binding site (27, 30).

**Fig. 2.**
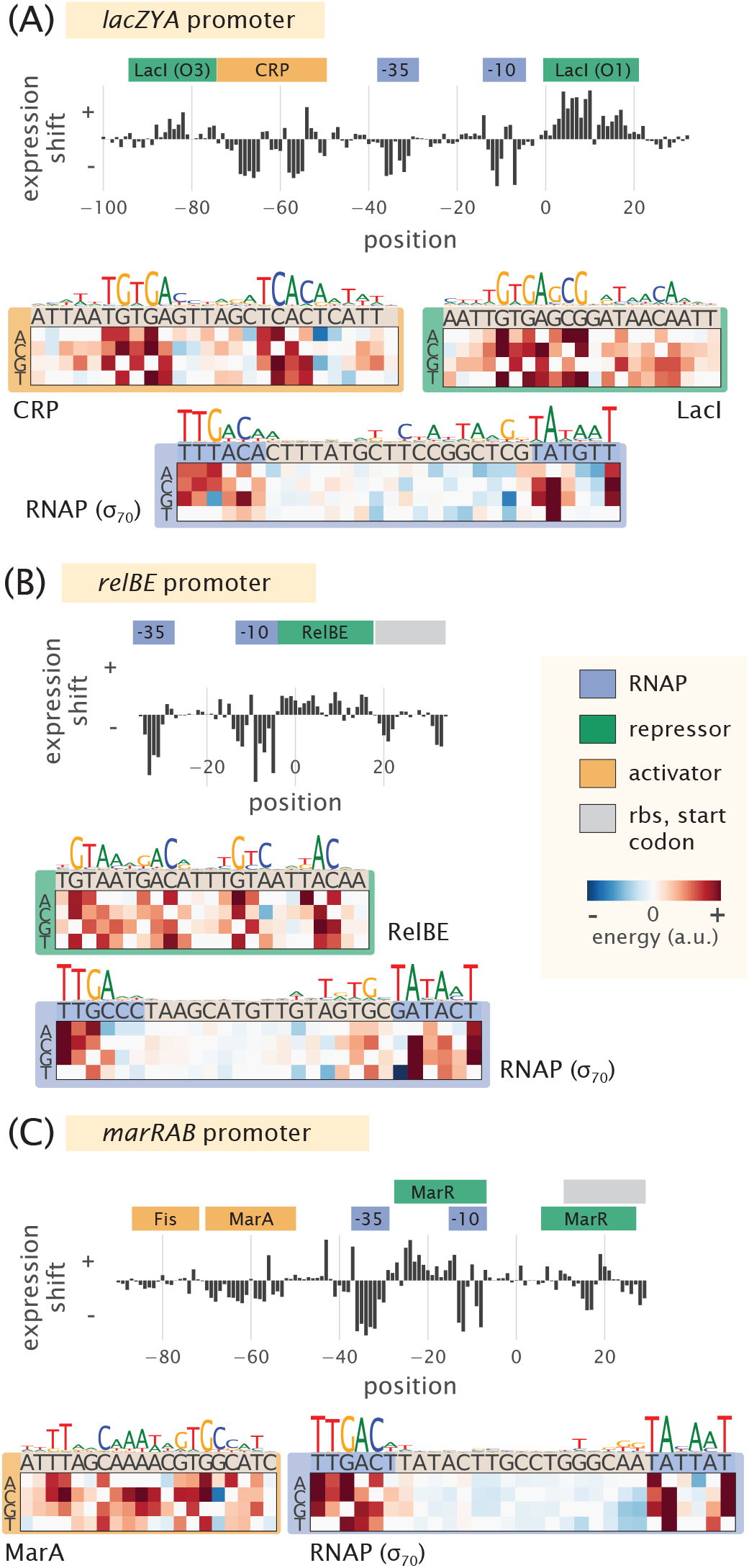
Characterization of the regulatory landscape of the *lac, rel*, and *mar* promoters. (A) Sort-Seq of the *lac* promoter. Cells were grown in M9 minimal media with 0.5% glucose at 37°C. Expression shifts are shown, with annotated binding sites for CRP (activator), RNAP (-10 and -35 subsites), and LacI (repressor) noted. Energy matrices and sequence logos are shown for each binding site. (B) Sort-Seq of the *rel* promoter. Cells were also grown in M9 minimal media with 0.5% glucose at 37°C. The expression shifts identify the binding sites of RNAP and RelBE (repressor), and energy matrices and sequence logos are shown for these. (C) Sort-Seq of the *mar* promoter. Here cells were grown in lysogeny broth (LB) at 30°C. The expression shifts identify the known binding sites of Fis and MarA (activators), RNAP, and MarR (repressor). Energy matrices and sequence logos are shown for MarA and RNAP. Annotated binding sites are based on those in RegulonDB.

Next we consider the *rel* promoter that transcribes the toxin-antitoxin pair RelE and RelB. It is one of about 36 toxin-antitoxin systems found on the chromosome, with important roles in cellular physiology including cellular persistence (31). When the toxin, RelE, is in excess of its cognate binding partner, the antitoxin RelB, the toxin causes cellular paralysis through cleavage of mRNA (32). Interestingly, the antitoxin protein also contains a DNA binding domain and is a repressor of its own promoter (33). We similarly performed Sort-Seq, with cells grown in M9 minimal media. The expression shifts are shown in Fig. 2B and were consistent with binding by RNAP and RelBE. In particular, a positive shift was observed at the binding site for RelBE, and the RNAP binding site mainly showed a negative shift in expression.

The third promoter, *mar*, is associated with multiple antibiotic resistance since its operon codes for the transcription factor MarA, which activates a variety of genes including the major multi-drug resistance efflux pump, ArcAB-tolC, and increases antibiotic tolerance (29). The *mar* promoter is itself activated by MarA, SoxS, and Rob (via the so-called marbox binding site), and further enhanced by Fis, which binds upstream of this marbox (34). Under standard laboratory growth it is under repression by MarR (29). We found that the promoter’s fluorescence was quite dim in M9 minimal media and instead grew libraries in lysogeny broth (LB) at 30° C (35). Again, the different features in the expression shift plot (Fig. 2C) appeared to be consistent with the noted binding sites. One exception was that the downstream MarR binding site was not especially apparent. Both positive and negative expression shifts were observed along its binding site, which may be due to overlap with other features present including the native ribosomal binding site. There have also been reported binding sites for CRP, Cra, CpxR/CpxA, and AcrR (1). However the studies associated with these annotations either required overexpression of the associated transcription factor, were computationally predicted, or demonstrated through *in vitro* assays and not necessarily expected under the growth condition considered here.

While each promoter qualitatively showed the expected regulatory behavior in each expression shift plot, it was important to show that we could also recover the quantitative features of binding by each transcription factor. Here we inferred energy matrices and associated sequence logos for the binding sites of RNAP, LacI, CRP, RelBE, MarA, and Fis. These are shown in Fig. 2A-C and Fig. S4, and indeed, agreed well with sequence logos generated from known genomic binding sites for these transcription factors (Pearson correlation coefficient r=0.5-0.9; see Supplemental Information Section C). For the repressors RelBE and MarR, there was no data available that characterized their sequence specificity with which to compare against. Here, instead, we validated our data by performing Sort-Seq in strains where the *relBE* or *marR* genes were deleted. In each case this resulted in a loss of the expression shift associated with binding by these repressors (Fig. 3), suggesting that the observed features are due to binding by these transcription factors.

**Fig. 3.**
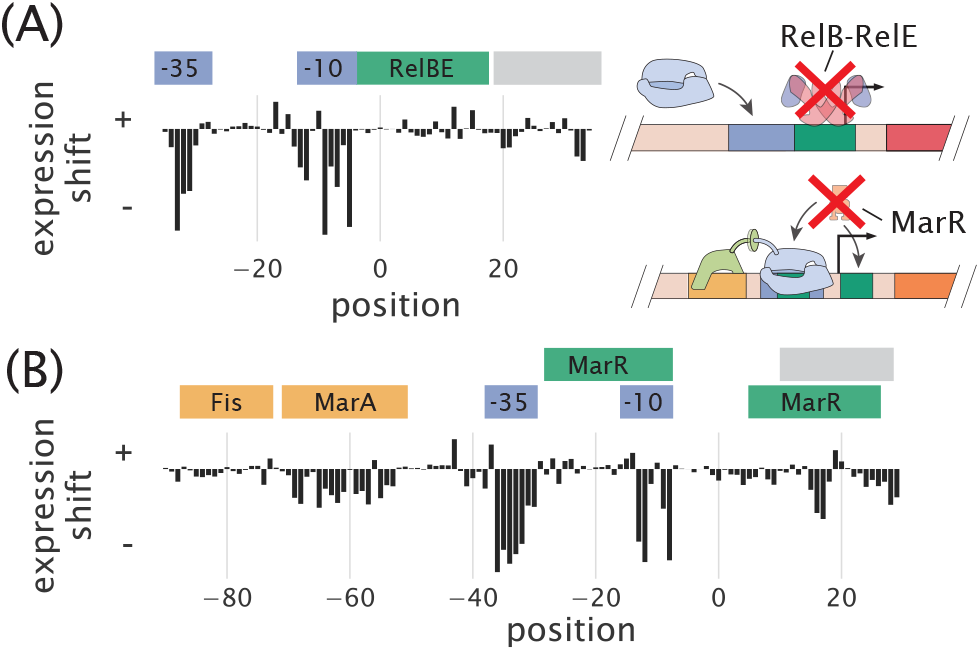
Expression shifts relfect binding by regulatory proteins. (A) Expression shifts for the *rel* promoter, but in a Δ*rel* genetic background. Cells were grown in conditions identical to Fig. 2B but do not show a positive expression shift across the entire RelBE binding site. (B) Expression shifts for the *mar* promoter, but in a Δ*marR* genetic background. The positive expression shift observed where MarR is expected to bind is no longer observed. Binding site annotations are identified in blue for RNAP sites, green for repressor sites, yellow for activator sites, and gray for ribosomal binding site and start codons. These annotations refer to the binding sites noted on RegulonDB that were observed in the Sort-Seq data.

### Identification of transcription factors with DNA affinity chromatography and quantitative mass spectrometry

It was next important to show that DNA affinity chromatography could be used to identify transcription factors in *E. coli*. In particular, a challenge arises in identifying transcription factors in most organisms due to their very low abundance. In *E. coli* the cumulative distribution in protein copy number shows that more than half have a copy number less than 100 per cell, with 90% having copy number less than 1,000 per cell. This is several orders of magnitude below that of many other cellular proteins (36).

We began by applying the approach to known binding sites for LacI and RelBE. For LacI, which is present in *E. coli* in about 10 copies per cell, we used the strongest binding site sequence, Oid (*in vivo K_d_* ~ 0.05 *nM*), and the weakest natural operator sequence, O3 (*in vivo K_d_* ≈ 110 *nM*) (27, 30, 37). In Fig. 4A we plot the protein enrichments from each transcription factor identified by mass spectrometry. LacI was found with both DNA targets, with fold enrichment greater than 10 in each case, and significantly higher than most of the proteins detected (indicated by the shaded region, which represents the 95% probability density region of all proteins detected, including non-DNA binding proteins). Purification of LacI with about 10 copies per cell using the weak O3 binding site sequence are near the limit of what would be necessary for most *E. coli* promoters.

**Fig. 4.**
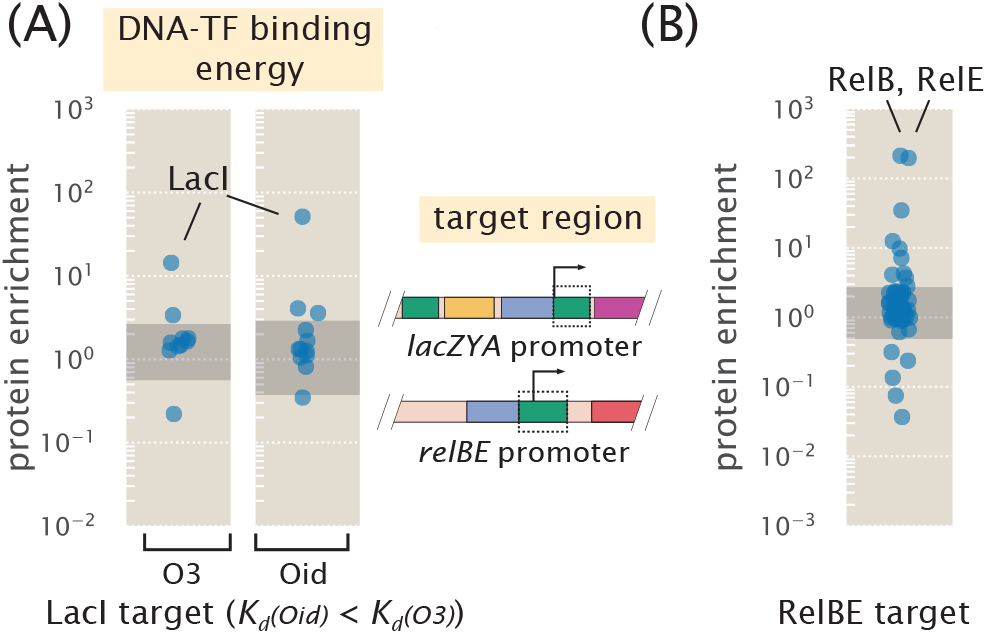
DNA affinity purification and identification of LacI and RelBE by mass spectrometry using known target binding sites. (A) Protein enrichment using the weak O3 binding site and strong synthetic Oid binding sites of LacI. LacI was the most significantly enriched protein in each purification. The target DNA region was based on the boxed area of the *lac* promoter schematic, but with the native O1 sequence replaced with either O3 or Oid. Data points represent average protein enrichment for each detected transcription factor, measured from a single purification experiment. (B) For purification using the RelBE binding site target, both RelB and its cognate binding partner RelE were significantly enriched. Data points show the average protein enrichment from two purification experiments. The target binding site is similarly shown by the boxed region of the *rel* promoter schematic. Data points in each purification show the protein enrichment for detected transcription factors. The gray shaded regions shows where 95% of all detected protein ratios were found.

To ensure this success was not specific to LacI, we also applied chromatography to the RelBE binding site. RelBE provides an interesting case since the strength of binding by RelB to DNA is dependent on whether RelE is bound in complex to RelB (with at least a 100 fold weaker dissociation constant reported in the absence of RelE (38, 39)). As shown in Fig. 4B, we found over 100 fold enrichment of both proteins by mass spectrometry. To provide some additional intuition into these results we also considered the predictions from a statistical mechanical model of DNA binding affinity (See Supplemental Information Section D). As a consequence of performing a second reference purification, we find that fold enrichment should mostly reflect the difference in binding energy between the DNA sequences used in the two purifications, and be much less dependent on whether the protein was in low or high abundance within the cell. This appeared to be the case when considering other *E. coli* strains with LacI copy numbers between about 10 and 1,000 copies per cell (Fig. S5C). Further characterization of the measurement sensitivity and dynamic range of this approach is noted in Supplemental Information Section E.

### Sort-Seq discovers regulatory architectures in unannotated regulatory regions

Given that more than half of the promoters in *E. coli* have no annotated transcription factor binding sites in RegulonDB, we narrowed our focus by using several high-throughput studies to identify candidate genes to apply our approach (40, 41). The work by Schmidt *et al.* (41) in particular measured the protein copy number of about half the *E. coli* genes across 22 distinct growth conditions. Using this data, we identified genes that had substantial differential gene expression patterns across growth conditions, thus hinting at the presence of regulation and even how that regulation is elicited by environmental conditions (see further details in Supplemental Information Section A and Fig. S2A-C). On the basis of this survey, we chose to investigate the promoters of *purT, xylE,* and *dgoRKADT.* To apply Sort-Seq in a more exploratory manner, we considered three 60 bp mutagenized windows spanning the intergenic region of each gene. While it is certainly possible that regulatory features will lie outside of this window, a search of known regulatory binding sites suggest that this should be sufficient to capture just over 70% of regulatory features in *E. coli* and provide a useful starting point (Fig. S6).

### The purT promoter contains a simple repression architecture and is repressed by PurR

The first of our candidate promoters is associated with expression of *purT*, one of two genes found in *E. coli* that catalyze the third step in *de novo* purine biosynthesis (42, 43). Due to a relatively short intergenic region, about 120 bp in length that is shared with a neighboring gene *yebG*, we also performed Sort-Seq on the *yebG* promoter (oriented in the opposite direction (44); see schematic in Fig. 5A). To begin our exploration of the *purT* and *yebG* promoters, we performed Sort-Seq with cells grown in M9 minimal media with 0.5% glucose. The associated expression shift plots are shown in Fig. 5A. While we performed Sort-Seq on a larger region than shown for each promoter, we only plot the regions where regulation was apparent.

**Fig. 5.**
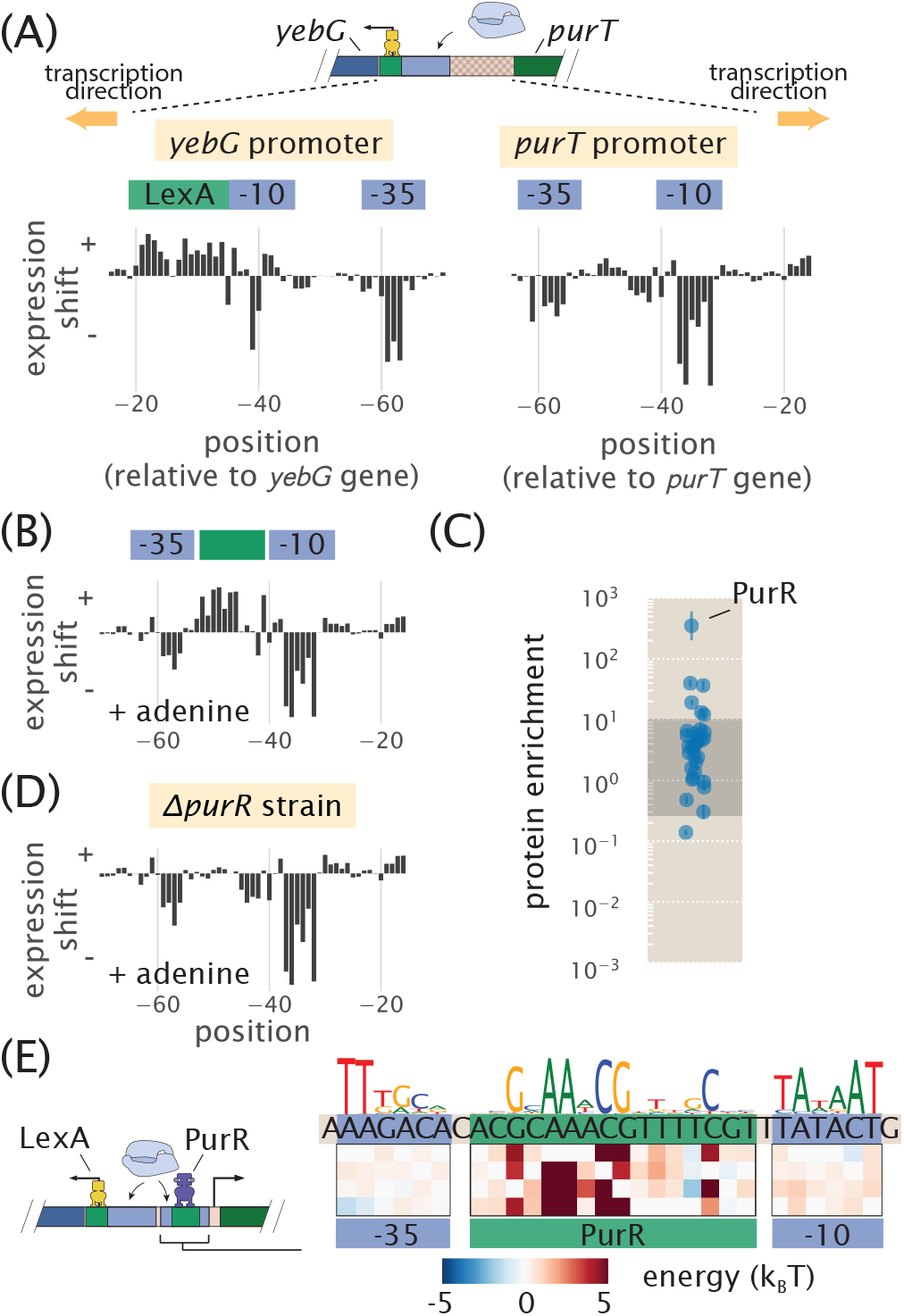
Sort-Seq distinguishes directional regulatory features and uncovers the regulatory architecture of the *purT* promoter. (A) A schematic is shown for the approximately 120 bp region between the *yebG* and *purT* genes, which code in opposite directions. Expression shifts are shown for 60 bp regions where regulation was observed for each promoter, with positions noted relative to the start codon of each native coding gene. Cells were grown in M9 minimal media with 0.5% glucose. The -10 and -35 RNAP binding sites of the *purT* promoter were determined through inference of an energy matrix and are identified in blue. (B) Expression shifts for the *purT* promoter, but in M9 minimal media with 0.5% glucose supplemented with adenine (100 *μ*g/ml). A putative repressor site is annotated in green. (C) DNA affinity chromatography was performed using the identified repressor site and protein enrichment values for transcription factors are plotted. Cell lysate was produced from cells grown in M9 minimal media with 0.5 % glucose. Binding was performed in the presence of hypoxanthine (10 *μ*g/ml). Error bars represent the standard error of the mean, calculated using log protein enrichment values from three replicates, and the gray shaded region represents 95% probability density region of all protein detected. (D) Identical to (B) but performed with cells containing a *ApurR* genetic background. (E) Summary of regulatory binding sites and transcription factors that bind within the intergenic region between the genes of *yebG* and *purT*. Energy weight matrices and sequence logos are shown for the PurR repressor and RNAP binding sites. Data was fit to a thermodynamic of simple repression, yielding energies in units of *k_B_T*.

For the *yebG* promoter, the features were largely consistent with prior work, containing a binding sites for LexA and RNAP. However, we found that the RNAP binding site is shifted 9 bp downstream from what was identified previously through a computational search (44), demonstrating the ability of our approach to identify and correct errors in the published record. We were also able to confirm that the *yebG* promoter was induced in response to DNA damage by repeating Sort-Seq in the presence of mitomycin C (a potent DNA cross-linker known to elicit the SOS response and proteolysis of LexA (45); see Fig. S7A, B, and D).

Given the role of *purT* in the synthesis of purines, and the tight control over purine concentrations within the cell (42), we performed Sort-Seq of the *purT* promoter in the presence or absence of the purine, adenine, in the growth media. In growth without adenine (Fig. 5A, right plot), we observed two negative regions in the expression shift plot. Through inference of an energy matrix, these two features were identified as the -10 and -35 regions of an RNAP binding site. While these two features were still present upon addition of adenine, as shown in Fig. 5B, this growth condition also revealed a putative repressor site between the -35 and -10 RNAP binding sites, indicated by a positive shift in expression (green annotation).

Following our strategy to find not only the regulatory sequences, but also their associated transcription factors, we next applied DNA affinity chromatography using this putative binding site sequence. In our initial attempt however, we were unable to identify any substantially enriched transcription factor (Fig. S7C). With repression observed only when cells were grown in the presence of adenine, we reasoned that the transcription factor may require a related ligand in order to bind the DNA, possibly through an allosteric mechanism. Importantly, we were able to infer an energy matrix to the putative repressor site whose sequence-specificity matched that of the well-characterized repressor, PurR (r=0.82; see Fig. S4). We also noted ChIP-chip data of PurR that suggests it might bind within this intergenic region (43). We therefore repeated the purification in the presence of hypoxanthine, which is a purine derivative that also binds PurR (46). As shown in Fig. 5C, we now observed a substantial enrichment of PurR with this putative binding site sequence. As further validation, we performed Sort-Seq once more in the adenine-rich growth condition, but in a Δ*purR* strain. In the absence of PurR, the putative repressor binding site disappeared (Fig. 5D), which is consistent with PurR binding at this location.

In Fig. 5E we summarize the regulatory features between the coding genes of *purT* and *yebG*, including the new features identified by Sort-Seq. With the appearance of a simple repression architecture (47) for the *purT* promoter, we extended our analysis by developing a thermodynamic model to describe repression by PurR. This enabled us to infer the binding energies of RNAP and PurR in absolute *k_B_T* energies (48), and we show the resulting model in Fig. 5E (see additional details in Supplemental Information Section Information H.3.4).

### The xylE operon is induced in the presence of xylose, mediated through binding of XylR and CRP

The next unannotated promoter we considered was associated with expression of *xylE,* a xylose/proton symporter involved in uptake of xylose. From our analysis of the Schmidt *et al.* (41) data, we found that *xylE* was sensitive to xylose and proceeded by performing Sort-Seq in cells grown in this carbon source. Interestingly, the promoter exhibited essentially no expression in other media (Fig. S7E). We were able to locate the RNAP binding site between -80 bp and -40 bp relative to the *xylE* gene (Fig. 6A, annotated in blue). In addition, the entire region upstream of the RNAP appeared to be involved in activating gene expression (annotated in orange in Fig. 6A), suggesting the possibility of multiple transcription factor binding sites.

**Fig. 6.**
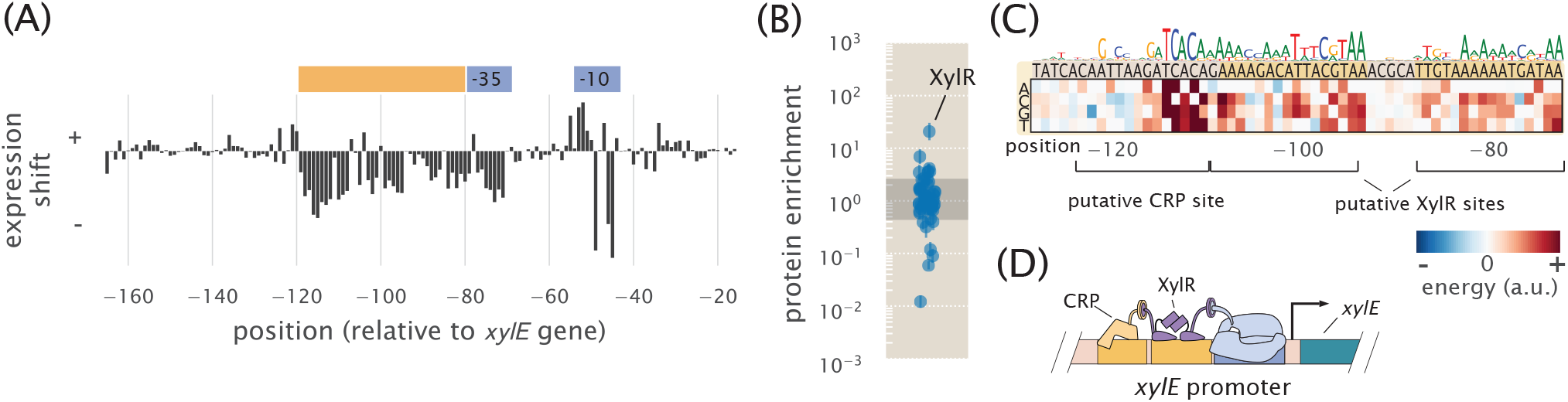
Sort-Seq identifies a set of activator binding sites that drive expression of RNAP at the *xylE* promoter. (A) Expression shifts are shown for the *xylE* promoter, with Sort-Seq performed on cells grown in M9 minimal media with 0.5% xylose. The -10 and -35 regions of an RNAP binding site (blue) and a putative activator region (orange) are annotated. (B) DNA affinity chromatography was performed using the putative activator region and protein enrichment values for transcription factors are plotted. Cell lysate was generated from cells grown in M9 minimal media with 0.5% xylose and binding was performed in the presence of xylose supplemented at the same concentration as during growth. Error bars represent the standard error of the mean, calculated using log protein enrichment values from three replicates. The gray shaded region represents 95% probability density region of all proteins detected. (C) An energy matrix was inferred for the region upstream of the RNAP binding site. The associated sequence logo is shown above the matrix. Two binding sites for XylR were identified (see also Fig. S4 and Fig. S7F) along with a CRP binding site. (D) Summary of regulatory features identified at *xylE* promoter, with the identification of an RNAP binding site and tandem binding sites for XylR and CRP.

We applied DNA affinity chromatography using a DNA target containing this entire upstream region. Due to the stringent requirement for xylose to be present for any measurable expression, xylose was supplemented in the lysate during binding with the target DNA. In Fig. 6B we plot the enrichment ratios from this purification and find XylR to be most significantly enriched. From an energy matrix inferred for the entire region upstream of the RNAP site, we were able to identify two correlated 15 bp regions (dark yellow shaded regions in Fig. 6C). Mutations of the XylR protein have been found to diminish transport of xylose (49), which in light of our result, may be due in part to a loss of activation and expression of this xylose/proton symporter. These binding sites were also similar to those found on two other promoters known to be regulated by XylR *(xylA* and *xylF* promoters), whose promoters also exhibit tandem XylR binding sites and strong binding energy predictions with our energy matrix (Fig. S7F).

Within the upstream activator region in Fig. 6A there still appeared to be a binding site unaccounted for with these tandem XylR binding sites. From the energy matrix, we were further able to identify a binding site for CRP, which is noted upstream of the XylR binding sites in Fig. 6C. While we did not observe a significant enrichment of CRP in our protein purification, the most energetically favorable sequence predicted by our model, TGCGACCNAGATCACA, closely matches the CRP consensus sequence of TGTGANNNNNNTCACA. In contrast to the *lac* promoter, binding by CRP here appears to depend more on the right half of the binding site sequence. CRP is known to activate promoters by multiple mechanisms (50), and CRP binding sites have been found adjacent to the activators XylR and AraC (49, 51), in line with our result. While further work will be needed to characterize the specific regulatory mechanism here, it appears that activation of RNAP is mediated by both CRP and XylR and we summarize this result in Fig. 6D (and considered further in Supplemental Information Section H.3.4).

### The dgoRKADT promoter is auto-repressed by DgoR, with transcription mediated by class II activation by CRP

As a final illustration of the approach developed here, we considered the unannotated promoter of *dgoRKADT.* The operon codes for D-galactonate-catabolizing enzymes; D-galactonate is a sugar acid that has been found as a product of galactose metabolism (52). We began by measuring expression from a non-mutagenized *dgoRKADT* promoter reporter to glucose, galactose, and D-galactonate. Cells grown in galactose exhibited higher expression than in glucose, as found by Schmidt *et al.* (41), and even higher expression when cells were grown in D-galactonate (Fig. S8A). This likely reflects the physiological role provided by the genes of this promoter, which appear necessary for metabolism of D-galactonate. We therefore proceeded by performing Sort-Seq with cells grown in either glucose or D-galactonate, since these appeared to represent distinct regulatory states, with expression low in glucose and high in D-galactonate. Expression shift plots from each growth conditions are shown in Fig. 7A.

**Fig. 7.**
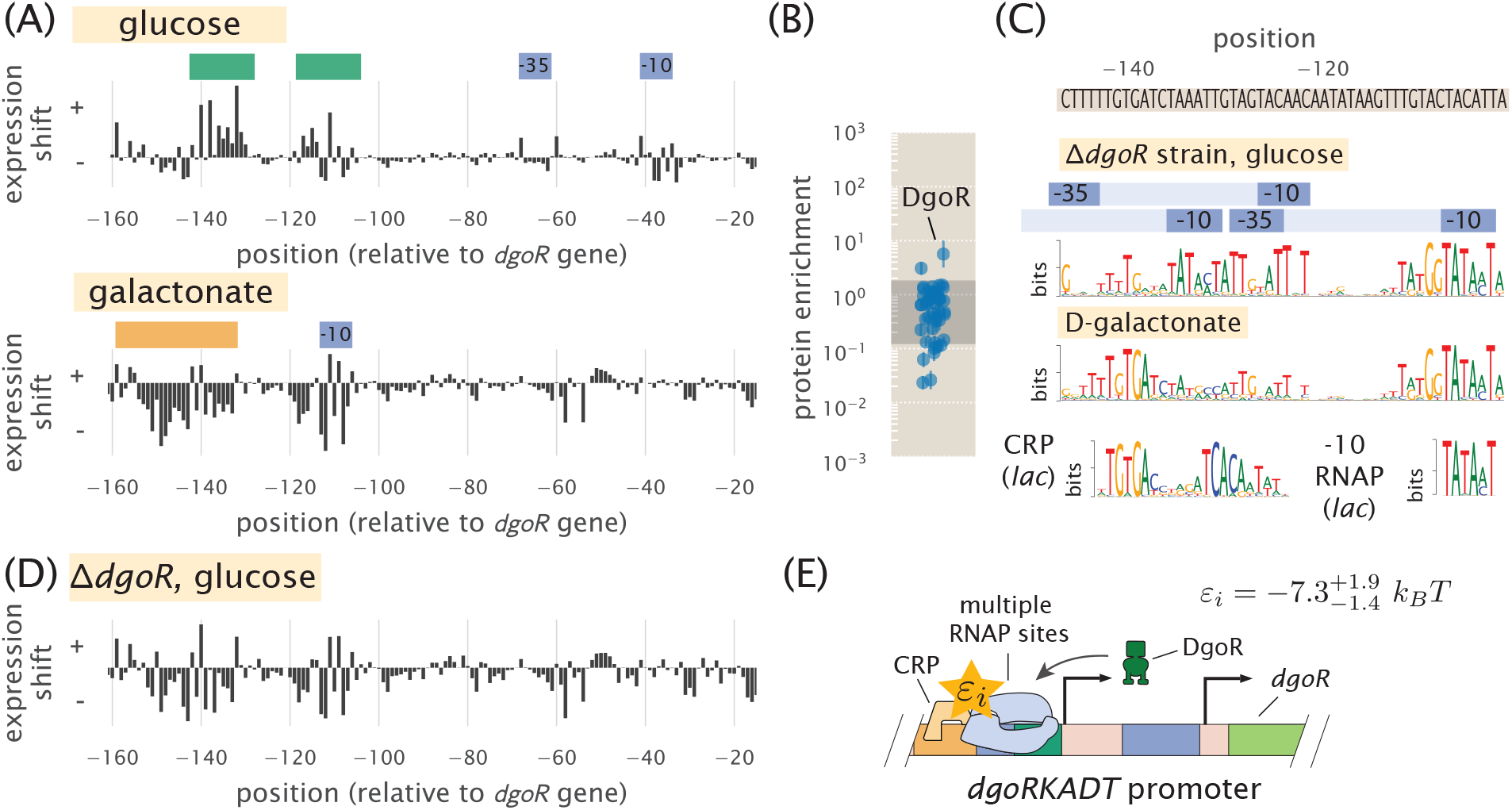
The *dgoRKADT* promoter is induced in the presence of D-galactonate due to loss of repression by DgoR and activation by CRP. (A) Expression shifts due to mutating the *dgoRKADT* promoter are shown for cells grown in M9 minimal media with either 0.5% glucose (top) or 0.23% D-galactonate (bottom). Regions identified as RNAP binding sites (-10 and -35) are shown in blue and putative activator and repressor binding sites are shown in orange and green, respectively. (B) DNA affinity purification was performed targeting the region between -145 to -110 of the *dgoRKADT* promoter. The transcription factor DgoR was found most enriched among the transcription factors plotted. Error bars represent the standard error of the mean, calculated using log protein enrichment values from three replicates, and the gray shaded region represents 95% probability density region of all proteins detected. (C) Sequence logos were inferred for the most upstream 60 bp region associated with the upstream RNAP binding site annotated in (A). Multiple RNAP binding sites were identified using Sort-Seq data performed in a Δ*dgoR* strain, grown in M9 minimal media with 0.5% glucose. (further detailed in Fig. S8). Below this, a sequence logo was also inferred using data from Sort-Seq performed on wild-type cells, grown in D-galactonate, identifying a CRP binding site (class II activation (50)). (D) Expression shifts are shown for the *dgoRKADT* promoter when performed in a Δ*dgoR* genetic background, grown in 0.5% glucose. This resembles growth in D-galactonate, suggesting D-galactonate may act as an inducer for DgoR. (E) Summary of regulatory features identified at *dgoRKADT* promoter, with the identification of multiple RNAP binding sites, and binding sites for DgoR and CRP. The interaction energy between CRP and RNAP, *εi*, was inferred to be 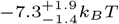, where the superscripts and subscripts represent the upper and lower bounds of the 95^th^ percentile of the parameter value distribution.

We begin by considering the results from growth in glucose (Fig. 7A, top plot). Here we identified an RNAP binding site between -30 bp and -70 bp, relative to the native start codon for *dgoR* (Fig. 7B). Another distinct feature was a positive expression shift in the region between -140 bp and -110 bp, suggesting the presence of a repressor binding site. Applying DNA affinity chromatography using this target region we observed an enrichment of DgoR (Fig. 7B), suggesting that the promoter is indeed under repression, and regulated by the first coding gene of its transcript. As further validation of binding by DgoR, the positive shift in expression was no longer observed when Sort-Seq was repeated in a Δ*dgoR* strain (Fig. 7D and Fig. S8C). We also were able to identify additional RNAP binding sites that were not apparent due to binding by DgoR. While only one RNAP -10 motif is clearly visible in the sequence logo shown Fig. 7C (top sequence logo; TATAAT consensus sequence), we used simulations to demonstrate that the entire sequence logo shown can be explained by the convolution of three overlapping RNAP binding sites (See Supplemental Information Section D and Fig. S8F).

Next we consider the D-galactonate growth condition (Fig. 7A, bottom plot). Like in the expression shift plot for the Δ*dgoR* strain grown in glucose, we no longer observe the positive expression shift between -140 bp and -110 bp. This suggests that DgoR may be induced by D-galactonate or a related metabolite. However, in comparison with the expression shifts in the Δ*dgoR* strain grown in glucose, there were some notable differences in the region between −160 bp and −140 bp. Here we find evidence for another CRP binding site. The sequence logo identifies the sequence TGTGA (Fig. 7C, bottom logo), which matches the left side of the CRP consensus sequence. In contrast to the *lac* and *xylE* promoters however, the right half of the binding site directly overlaps with where we would expect to find a -35 RNAP binding site. This type of interaction by CRP has been previously observed and is defined as class II CRP dependent activation (50), though this sequence-specificity has not been previously described.

In order to isolate and better identify this putative CRP binding site we repeated Sort-Seq in *E. coli* strain JK10, grown in 500 *μ*M cAMP. Strain JK10 lacks adenlyate cyclase *(cyaA)* and phosphodiesterase (*cpdA*), which are needed for cAMP synthesis and degradation, respectively, and is thus unable to control intracellular cAMP levels necessary for activation by CRP (derivative of TK310 (37)). Growth in the presence of 500 *μ*M cAMP provided strong induction from the *dgoRKADT* promoter and resulted in a sequence logo at the putative CRP binding site that even more clearly resembled binding by CRP (Fig. S8E). This is likely because expression is now dominated by the CRP activated RNAP binding site. Importantly, this data allowed us to further infer the interaction energy between CRP and RNAP, which we estimate to be -7.3 *k_B_T* (further detailed in Supplemental Information Section H.3.4). We summarize the identified regulatory features in Fig. 7E.

## Discussion

We have established a systematic procedure for dissecting the functional mechanisms of previously uncharacterized regulatory sequences in bacteria. A massively parallel reporter assay, Sort-Seq (12), is used to first elucidate the locations of functional transcription factor binding sites. DNA oligonucleotides containing these binding sites are then used to enrich the cognate transcription factors and identify them by mass spectrometry analysis. Information-based modeling and inference of energy matrices that describe the DNA binding specificity of regulatory factors provide further quantitative insight into transcription factor identity and the growth condition dependent regulatory architectures.

To validate this approach we examined four previously annotated promoters of *lac*, *rel*, *mar*, and *yebG*, with our results consistent with established knowledge (12, 27, 29, 30, 35, 39). For the *yebG* promoter, however, our approach corrected an error in a previous annotation. Importantly, we find that DNA affinity chromatography experiments on these promoters were highly sensitive. In particular, LacI was unambiguously identified with the weak O3 binding site, even though LacI is present in only about 10 copies per cell (30). Emboldened by this success, we then studied promoters having little or no prior regulatory annotation: *purT, xylE,* and *dgoR.* Here our analysis led to a collection of new regulatory hypotheses. For the *purT* promoter, we identified a simple repression architecture (47), with repression by PurR. The *xylE* promoter was found to undergo activation only when cells are grown in xylose, likely due to allosteric interaction between the activator XylR and xylose, and activation by CRP (49, 51). Finally, in the case of *dgoR*, the base-pair resolution allowed us to tease apart overlapping regulatory binding sites, identify multiple RNAP binding sites along the length of the promoter, and infer further quantitative detail about the interaction between the newly identified binding sites for CRP and RNAP. We view these results as a critical first step in the quantitative dissection of transcriptional regulation, which will ultimately be needed for a predictive understanding of how such regulation works.

An important aspect of the presented approach is that it is readily parallelized and scalable. There are a number of ways to increase the resolution and throughput. Microarray-synthesized promoter libraries should allow multiple loci to be studied simultaneously. Landing pad technologies for chromosomal integration (53) should enable massively parallel reporter assays to be performed in chromosomes instead of on plasmids. Techniques that combine these assays with transcription start site readout (54) may further allow the molecular regulators of overlapping RNAP binding sites to be deconvolved, or the contributions from separate RNAP binding sites, like those observed on the *dgoR* promoter, to be better distinguished. Although our work was directed toward regulatory regions of *E. coli*, there are no intrinsic limitations that restrict the analysis to this organism. Rather, it should be applicable to any bacterium that supports efficient transformation by plasmids. And although we have focused on bacteria, our general strategy should be feasible in a number of eukaryotic systems – including human cell culture – using massively parallel reporter assays (13–15) and DNA-mediated protein pull-down methods (20, 21) that have already been established.

## Materials and Methods

See Supplemental Information Section I for extended experimental details.

### Bacterial strains

All *E. coli* strains used in this work were derived from K-12 MG1655, with deletion strains generated by the lambda red recombinase method (55). In the case of deletions for *lysA* (Δ*lysA*::*kan*), *purR* (Δ*purR*::*kan*), and *xylE* (Δ*xylE*::*kan*), strains were obtained from the Coli Genetic Stock Center (CGSC, Yale University, CT, USA) and transferred into a fresh MG1655 strain using P1 transduction. The others were generated in house and include the following deletion strains: Δ*lαcIZY A*, Δ*relBE*::*kan*, Δ*marR*::kan, Δ*dgoR*::kan (see Supplemental Information Section I.1 for details on strain construction).

### Sort-Seq

Mutagenized single-stranded oligonucleotide pools were purchased from Integrated DNA Technologies (Coralville, IA), with a target mutation rate of 9%. Note that in the case of the *lacZ* promoter, the library is identical to that used in the experiments of Razo-Mejia *et al.* (56), and had a mutation rate of approximately 3%. Library oligonucleotides were PCR amplified and inserted into the PCR amplified plasmid backbone (i.e. vector) of pJK14 (SC101 origin) (12) by Gibson assembly and electroporated into cells following drop dialysis in water.

Cells were grown to saturation in LB and then diluted 1:10,000 into the appropriate growth media for the promoter under consideration. Upon reaching an OD600 of about 0.3, the cells were washed two times with chilled PBS by spinning down the cells at 4000 rpm for 10 minutes at 4°C and diluted to an OD of 0.1-0.15. A Beckman Coulter MoFlo XDP cell sorter was used to sort cells by fluorescence, with 500,000 cells collected into each of the four bins. Sorted cells were then re-grown overnight in 10 ml of LB media, under kanamycin selection. The plasmid in each bin were miniprepped following overnight growth (Qiagen, Germany) and PCR was used to amplify the mutated region from each plasmid for Illumina sequencing (see Supplemental Information Section I.3 and I.4 for additional Sort-Seq and sequencing details, respectively). Details on constructing expression shift plots and the model inference that was performed are provided in Supplemental Information Section H.

### DNA affinity chromatography

SILAC labeling (26) was implemented by growing cells in either the stable isotopic form of lysine (^13^C_6_H_14_^15^N_2_O_2_), referred to as the heavy label, or natural lysine, referred to as the light label. Cell lysates were prepared using Δ*lysA* cells. For each heavy and light labelled cells, 500 ml M9 minimal media was inoculated 1:5,000 with an overnight LB culture of Δ*lysA* cells, and grown to an OD600 of ~ 0.6 (supplemented with the appropriate lysine; 40 *μ*g/ml). Cultures were pelleted, lyse using a Cell Disruptor (CF Range, Constant Systems Ltd., UK) and concentrated to ~150 mg/ml using Amicon Ultra-15 centrifugation units (3kDa MWCO, Millipore).

DNA affinity chromatography was performed by incubating cell lysate with magnetic beads (Dynabeads MyOne T1, ThermoFisher, Waltham, MA) containing tethered DNA. The DNA was tethered through a linkage between streptavidin on the beads and biotin on the DNA. Single-stranded DNA was purchased from Integrated DNA Technologies with the biotin modification on the 5’ end of the oligonucleotide sense strand. Cell lysates were incubated on a rotating wheel with the DNA tethered beads overnight at 4°C. Beads were washed three times using lysis buffer and once more with NEB Buffer 3.1 (New England Biolabs, MA, USA). Both purifications (with the target DNA and reference control) were combined by resuspending in 50 *μ*L NEB Buffer 3.1, and the DNA was cleaved by adding 10 *μ*l of the restriction enzyme PstI (100,000 units/ml, New England Biolabs targeting a CTGCAG sequence on the DNA) and incubating for 1.5 hours at 25°C. The beads were then removed and the samples prepared for mass spectrometry by in-gel digestion with endoproteinase Lys-C.

### LC-MS/MS analysis and protein quantitation

Liquid chromatography tandem-mass spectrometry (LC-MS/MS) experiments were carried out as previously described (57) and further detailed in supplemental experimental details. Thermo RAW files were processed using MaxQuant (v. 1.5.3.30) (58). Spectra were searched against the UniProt *E. coli* K-12 database (4318 sequences) as well as a contaminant database (256 sequences). Additional details are provided in Supplemental Information Section I.5. To calculate the overall protein ratio, the non-normalized protein replicate ratios were log transformed and then shifted so that the median protein log ratio within each replicate was zero (i.e., the median protein ratio was 1:1). The overall experimental log ratio was then calculated from the average of the replicate ratios.

### Code and data availability

All code used for processing data and plotting, as well as the final processed data are available upon request. Thermo RAW files for mass spectrometry are available on the jPOSTrepo repository (59) under accession code PXD007892. Sort-Seq sequencing files are available on the Sequence Read Archive under accession code SRP121362.

## Acknowledgements

We thank David Tirrell, Bradley Silverman, and Seth Lieblich for access and training for use of their Beckman Coulter MoFlo XDP cell sorter. We thank Jost Vielmetter and Nina Budaeva for access and training for use on their Cell Disruptor. We also thank Hernan Garcia, Manuel Razo-Mejia, Griffin Chure, Suzannah Beeler, Heun Jin Lee, Justin Bois, and Soichi Hirokawa for useful advice and discussion. This work was supported by La Fondation Pierre-Gilles de Gennes, the Rosen Center at Caltech, and the National Institutes of Health DP1 OD000217 (Director’s Pioneer Award), R01 GM085286, and 1R35 GM118043-01 (MIRA), the Gordon and Betty Moore Foundation through GBMF227, the National Institutes of Health 1S10RR029594-01A1 and the Beckman Institute. NB is an HHMI International Student Research fellow.

